# Temporal experience modifies future thoughts: Manipulation of Libet’s W influences difficulty assessment during a decision-making task

**DOI:** 10.1101/2020.08.03.233940

**Authors:** Eve A. Isham

## Abstract

Past studies have employed the subjective experience of decision time (Libet’s W) as an index of consciousness, marking the moment at which the agent first becomes aware of a decision. In the current study, we examined whether the temporal experience of W affects subsequent experience related to the action. Specifically, we tested whether W influenced the perception of difficulty in a decision-making task, hypothesizing that temporal awareness of W might influence the sense of difficulty. Consistent with our predictions, when W was perceived as early or late, participants subsequently rated the decision difficulty to be easy or difficult, respectively (Exp.1). Further investigation showed that perceived difficulty, however, did not influence W (Exp.2). Together, our findings suggest a unidirectional relationship such that W plays a role in the metacognition of difficulty evaluation. The results imply that subjective temporal experience of decision time modifies the consequential sense of difficulty.

**Highlights:**

- Perceived timing of decision (W) can bias the metacognition of difficulty evaluation in a decision-making task.
- Defined as a temporal index of consciousness, time W’s influence on difficulty evaluation reflects the possibility that the role of consciousness is to modify subsequent thoughts and behaviors.
- Explicit attention is necessary for the timing of decision (W) to be consciously experienced and effectively influential on subsequent thoughts.

---

A seminal study by Libet and colleagues examined the timing of decisions, questioning whether volition played a causal role in action output (Libet, Gleason, Wright and Pearl, 1983). In the paradigm, participants performed a simple voluntary act (e.g., wrist flexion or a button press) and reported the time they felt the urge to act. This reported time, termed W, is often taken to signify the earliest moment in which one becomes aware of the moment of decision to execute an action. When evaluated in the context of a neural signature, termed the readiness potential (Kornhuber & Deecke, 1965), Libet observed that this brain activity preceded participant reports of W by approximately 300-800 ms (Libet et al., 1983). Importantly, the latency of W relative to the timing of the readiness potential led to the interpretation that a conscious decision to act does not actually cause an action. In support of this view, Masicampo and Baumeister (2013) explained that the latency of W occurred too late in the event to be involved in the initiation of behaviors. This led to the controversial interpretation that free will is simply an illusion (Wegner, 2002), further prompting an unsettled debate of whether conscious experiences, at least as measured by the subjective assessment of W, have a function at all.

While Libet’s data led to the interpretation that consciousness has no function, Libet (1985) argued that in fact it did by pointing out that conscious will could have an influence on the action outcome in other circumstances. He postulated that consciousness alerts the system to reconsider, and possibly cancel, unconsciously-initiated actions (Libet, 1985, 2011). Libet proposed that such a cancelation process occurs during the temporal window between W and the voluntary motor response (e.g., a keypress). Although the presence of the readiness potential prior to judgments of action initiation might appear to argue for deterministic action, the ability to modify or cancel actions after a decision is reached could still serve as evidence for functional and purposeful consciousness.

A number of other theoretical proposals have suggested that consciousness serves the purpose of influencing and modifying future thoughts and behaviors. For instance, Masicampo and Baumeister proposed that conscious thought may have indirect effects on behavior, even if it does not directly control it (Baumeister, Masicampo, & Vohs, 2011; Masicampo & Baumeister, 2013). For example, a study on mental simulations showed that experimentally enhancing the link between thought and action led to better performance on an exam in college students (Pham & Taylor, 1999). This perspective thus implies that conscious thoughts and mental content aid in the modification and planning of future behaviors (Pockett, 2004; Wegner, 2002).

In a recent study, we measured W and perception of difficulty during a decision-making task (Isham, Wulf, Mejia, & Krisst, 2017). We found that judgments of W correlated with difficulty ratings such that later judgments of W corresponded with higher difficulty ratings and earlier judgments of W corresponded with lower difficulty ratings. In this previous study, we speculated that the temporal experience of W might influence the metacognition of difficulty assessment, although we did not directly test this issue. This observation gave rise to the current research question of whether the awareness of time W causally impacts successive thoughts and ideas.

If W can, in some instances, causally influence difficulty ratings, such findings might suggest that consciousness (as indexed by W) has a role in modifying subsequent thoughts and behaviors. A framework that explains this possibility is the Attended Intermediate-Level Representation theory (AIR) offered by Prinz (2000). According to AIR theory, consciousness serves the function of sending viewpoint specific information into the intermediate-level of processing (i.e., working memory). In turn, this allows for sequential planning and decisions throughout the course of action (Prinz, 2005). With respect to temporal consciousness in the Libet paradigm, it is possible that the timing of decision (i.e., W), which typically resides in the lower sensory level (e.g., Mylopoulous, 2015), is brought forth to the intermediate, conscious level when the participants are asked to report W. As Prinz describes, at this intermediate level, information is represented in a specific viewpoint. Relating this to the results from Isham et al. (2017), an awareness of early or late W could therefore have an influence on such a viewpoint, thereby influencing metacognition regarding difficulty evaluation.

In contrast to the above speculation, one could argue that the relationship between W and difficulty assessment observed in the Isham et al. study (2017) is correlational rather than causal. Moreover, this relationship may be mediated by response time given it has been shown that metacognition of task performance (e.g., confidence ratings) depends partially on how fast one completes the task (e.g., Fleming, Massoni, Gajdos, & Vergnaud, 2016). In Isham et al., participants had a sense of their response time (i.e., early vs late); in this manner, we cannot rule out the possible influence of response time on difficulty ratings from this earlier study. To successfully determine the influence of W on difficulty ratings, we must also rule out this simpler explanation of a mediating influence of response time on both W and difficulty ratings.

In the current study, we aimed to 1) demonstrate that W, rather than response time, has an influence on difficulty ratings (Experiment 1); and 2) establish a causal relationship between W and difficulty ratings (Experiments 1 and 2). To meet Aim 1, we compared the effects of W and response time on difficulty ratings. We set up this assay experimentally such that in one condition, participants reported W, while in another condition, participants made no report of W at all (Experiment 1). A sense of how long it took to decide (i.e., response time), however, was present in both cases. If W were to influence difficulty judgment, independent of response time, then we might expect to see an effect of W only in the condition that required a W report. On the other hand, if response time were the primary contributor to the fluctuation in difficulty judgment, then the difficulty ratings would vary with response time despite the presence or absence of the W report.

Upon establishing that W has an influence on difficulty ratings in Experiment 1, we next examined whether the observed effect is a reflection of a causal relationship between W and difficulty ratings. We conducted Experiment 2 to determine whether W would vary with a difficulty rating manipulation. If the difficulty manipulation influenced the W report, then the relationship would be bidirectional and therefore not causal. On the other hand, if the manipulated difficulty had no influence on the W report, it would imply that W influences difficulty judgment, but not the reverse. Such a causal relationship, if established by our study, would imply that temporal awareness, as indexed by W, has an influential role in subsequent thoughts and ideas.

## EXPERIMENT 1

Participants performed a binary decision-making task by indicating whether they agreed or disagreed with different scenarios presented to them. They also reported or omitted time W, and rated difficulty of each decision made. The goal of Experiment 1 was to demonstrate that W, but not response time, had an influence on difficulty assessment in a decision making task. To achieve this goal, we biased the participants’ perception of W to be earlier or later via a tone manipulation procedure (Banks & Isham, 2009). We anticipated the difficulty ratings to vary with W but not with response time. All data and stimulus materials are available upon request.

### Method

#### Participants

Forty-eight participants were recruited from a pool of undergraduate student volunteers (38 females, 18-35 years old) at the University of California, Davis. All participants were fluent in English. The participants consented in writing to the study protocol which followed the guidelines approved by the Institutional Review Board of the University of California, Davis.

#### Rationale and estimation of sample size

The sample size estimation was computed based on our previous study (Isham et al., 2017), which observed the effect size of W associated with easy and difficult to be .31. Based on this, a power analysis revealed a sample size of 23 to be sufficient at alpha = .05, and power = 0.8 (G*Power 3, Faul, Erdfelder, Lang, & Buchner, 2007) for the W group. The sample size was increased to 24 in the current study in anticipation of outliers.

For response time, we anticipated a small effect in the two comparisons of interest, one of which was a between-subjects variable (W Report vs No W Report Group). Therefore, we used an estimated effect size of .2. The estimated sample size was 40, suggesting that approximately 20 participants were needed in each of the between-subjects conditions. Given that 24 participants were needed to in the W Report Group as previously described, we therefore recruited 24 participants for the No W Report Group, resulting in 48 participants total for Experiment 1.

#### Stimuli and Apparatus

In Experiment 1, the participants listened to audio statements and were instructed to indicate whether they agreed or disagreed with each statement. Next they reported time W by reading the time from an analogue clock (like that used in Libet et al., 1983). Then, they rated the difficulty of the decision (summarized in Figure 1 and described in greater detail in the Procedure section below). In one experimental condition (No W Report condition), participants also answered a questionnaire at the conclusion of the experiment. To observe the influence of W on the difficulty ratings, we also manipulated the subjective report of W using a tone-manipulation procedure (Banks & Isham, 2009). The statement stimuli, Libet clock, the tone used in the manipulation, and the post-interview questions are described below.

**Figure 1.**
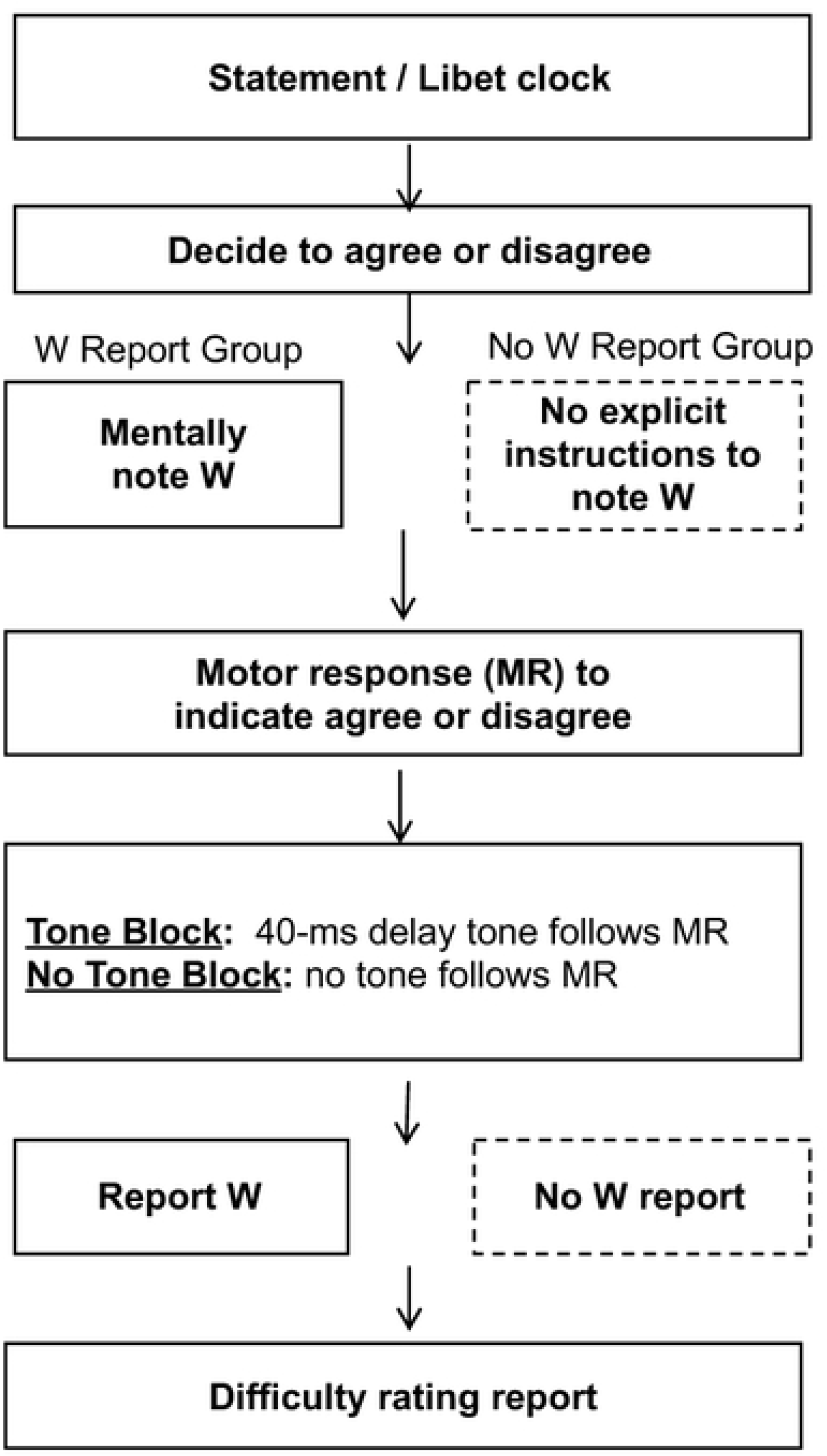
Experiment 1 procedure.

##### Statement stimuli

There were 111 statements total (see sample stimuli in Appendix 1; adapted from Isham et al., 2017). The statements had varying degrees of consequences which presumably would render different degrees of decisional difficulty. For example, an agree/disagree decision to the statement “I like red more than blue” was assumed to have less consequence and thus lower difficulty than a decision to the statement “To save a village, it’s okay to sacrifice a child.” The statements were recorded using a female speaker. The audio statements were edited using the software Audacity (Audacity, Boston, MA). The average statement length was 3037 ms (*SD* = 912). The statements were randomized and serially presented using Superlab (Cedrus Corporation, San Pedro, CA).

##### Libet Clock

While each statement was presented auditorily via the computer speakers, an analogue circular clock was simultaneously presented visually on the computer monitor, sharing the same onset time as the statement stimulus. The clock was 10.16 cm in diameter, positioned at the center of the screen, and was viewed from approximately 60 cm. Sixty tick marks were drawn along the circumference; a red dot moved along these tick marks, completing a rotation in 3 seconds. Each trial concluded at the end of the fourth clock rotation, giving each trial a duration of 12 seconds. The clock did not stop in response to the button press.

##### Tone

A critical component of the current experiment was to manipulate the perception of W to be later using a tone manipulation procedure (Banks & Isham, 2009). The tone was created from a sine wave at 1000 Hz frequency and approximately 50 ms in duration (Audacity, Boston MA). As described in the Procedure section below, in half of the trials, the tone was delivered 40 ms after the decision was made (i.e., a keypress) while the remaining trials did not receive a tone.

Our prior work has shown that a tone, when presented after a brief delay, shifts time W to be later compared to when there is no tone (Banks & Isham, 2009). From this earlier study, we used 5, 20, 40 and 60 ms delay tone (the 5 ms, rather than 0 ms, was the shortest that our technology was able to implement). Along with unpublished data in from our lab, we observed W to be earliest when the tone was absent, with W shifting systematically to be later with the occurrence of the tone, demonstrating a form of temporal binding. Because a primary interest was to manipulate participants’ subjective W to be as early or as late as possible, we decided to use no tone and a 40 ms delay tone to maximize the effect. A 40 ms delay tone was more favorable than the 60 ms delay tone because temporal binding has been shown to decline around 60-80 ms (Eagleman & Sejnowski, 2007)

##### Post-experiment Interview

Participants in the No W Report group received a two-item post-experiment questionnaire to ensure that the time component was not explicitly considered.

1. *“In terms of time, please elaborate on what you were thinking of when watching the clock.”* This question was used to assess whether participants explicitly thought of their decision time. No participants reported using the clock to time their decision and the clock did not prompt them to think about the timing of their decisions.
2. *“Please elaborate on your experience with the tone.”* This question was used to assess whether the participants were aware of the delay between timing of keypress and the tone presentation. No participants reported awareness of the delay manipulation.

#### Procedure

Figure 1 illustrates the step-by-step procedure in Experiment 1. The participants were randomly assigned to either the W Report or No W Report group. In both groups, the participants were instructed to listen to each statement carefully and to make a decision to agree or disagree with the statement. While listening to each statement, the participants were also viewing the Libet clock. Those who were assigned to the W Report group were instructed to make a mental note of the position of the red dot on the clock when they felt they had reached a decision (i.e. “W”). It was emphasized that this was not the time in which they physically pressed the button but rather the earliest moment in which they became aware of having an inkling toward a decision. Participants in the No W Report group were also instructed to watch the clock but did not receive explicit instructions to note time W.

When a decision was reached, the participants indicated their decision by pressing the AGREE or DISAGREE key on the keyboard (we alternated between the A and the L keys). This motor response elicited a brief tone, delivered 40 ms after the keypress in half of the trials (Tone Block). In the remaining half of the trials (No Tone Block), no tone was delivered upon keypress. The order of Tone and No Tone blocks were randomized across participants.

Upon the motor response and tone delivery, when applicable, the participants in the W Report group verbally reported the time of their decision that was read from the clock, followed by a difficulty rating on ta scale of 1 to 5, with 5 being the most difficult. Participants in the No W Report group reported only the difficulty rating and answered the two-item questionnaire about their temporal experience.

A total of 10 practice trials and 111 experimental trials were administered to each participant. Of these, half were randomly assigned to a Tone block and the remaining trials were randomly assigned to the No Tone block. The order of the Tone and No Tone blocks was counterbalanced. Note that the first of the two blocks always consisted of 56 trials; the additional trial served as an extra practice trial.

#### Analysis

The design of the experiment consisted of three independent variables: W Report Group (W is reported or W is not reported), Tone Presence (Delayed Tone or No Tone), and Time Course (First half of the experiment or Block 1, and Second half of the experiment or Block 2). Three dependent variables were measured: W time, response time, and difficulty ratings. Response time was measured from the moment the statement ended to the time of keypress; greater response time indicates that the keypress was made later. W, as traditionally computed, was backward-referenced from the moment of action. A greater W value indicates that the perceived moment of decision was earlier and was further away from the time of action.

##### Bayes Factor Analysis and False Discovery Rate Correction

Bayes Factor analysis for all t-tests was included (Rouder, Speckman, Sun, Morey, & Iverson, 2009). For results below the significance threshold (p < 0.05), we used a Bayes Factor, BF_10_, to indicate the degree of favorability toward the alternative hypothesis. For results that were not below the significance threshold, the Bayes Null factor, BF_01_, was used. The Cauchy prior r-scale was set at the default value of d = .707. The larger the Bayes Factor, the greater the evidence in support of the corresponding hypothesis being tested.

To minimize the false discovery rate, we employed the Benjamini-Hochberg procedure for multiple comparison corrections (Benjamini, Drai, Elmer, Kafkafi, & Golani, 2001).

### Results

#### W vs Response Time

To determine that W had an influence on difficulty ratings, we must rule out a simpler explanation of a mediating influence of response time on both W and difficulty ratings. To do so, we first must establish a scenario in which W and response time behave differently. According to our hypotheses, the presence or absence of the delayed post-action tone manipulation would bias the W report but not response time. The results are as follows:

##### W time

Time of W was measured as the timing of decision relative to the timing of action. To illustrate the effect of the tone manipulation on W, the W reports were subjected to a paired sample t-test comparing the Delayed Tone and No Tone conditions. Replicating our previous work (Banks & Isham, 2009), we observed the tone manipulation successfully altered the perceived time of decision such that when a delayed tone was given, the perceived time of decision (*M* = 371.31 ms before keypress, *SE* = 38.60) was later than when no tone was given (*M* = 410.90 ms before keypress, *SE* = 38.49), *t*(23) = 2.47, *p* = .021, corrected for multiple comparisons (BH *p*- value = .042), BF_10_ = 2.58. The mean W times are represented in Figure 2A.

**Figure 2.**
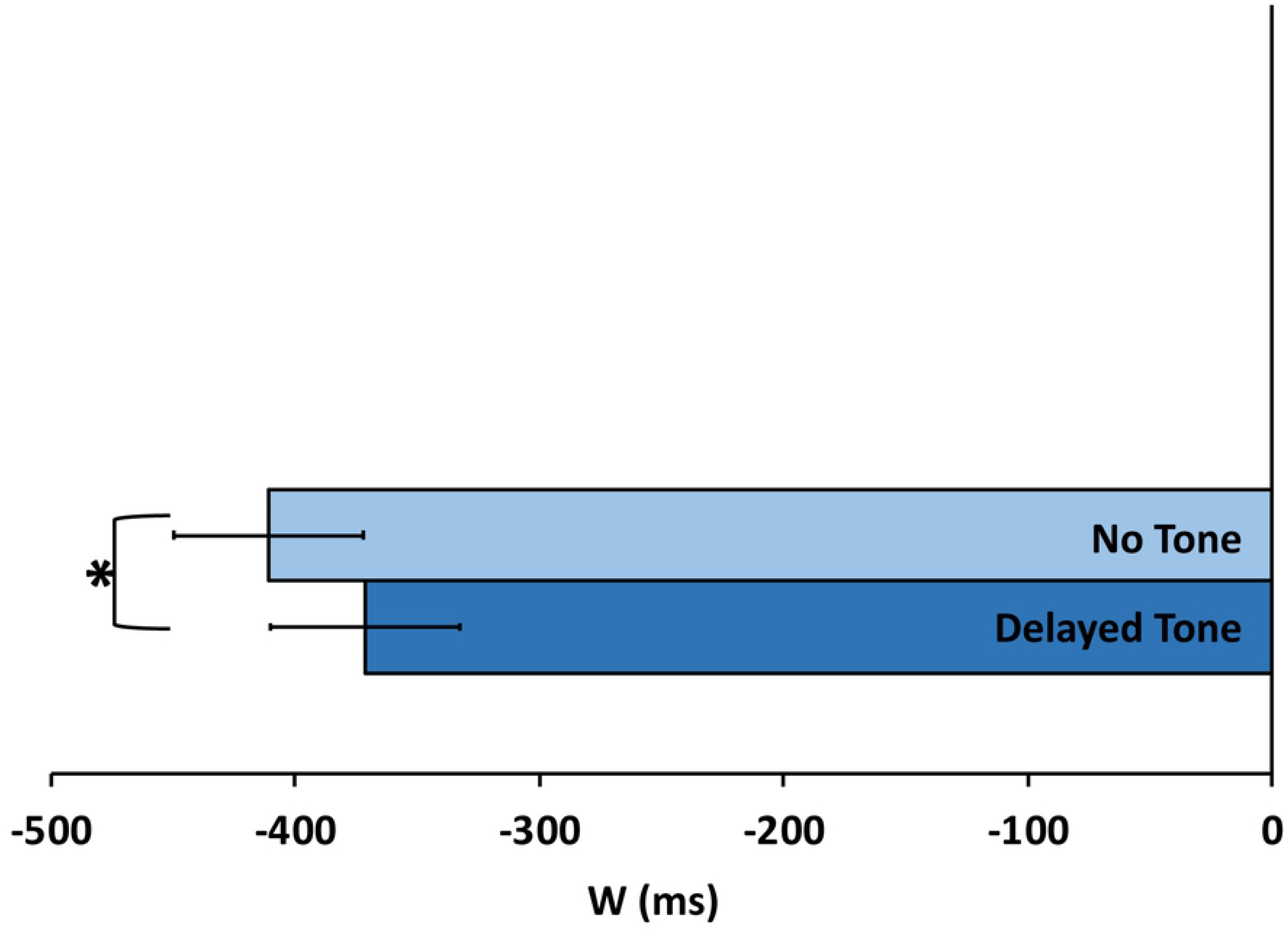
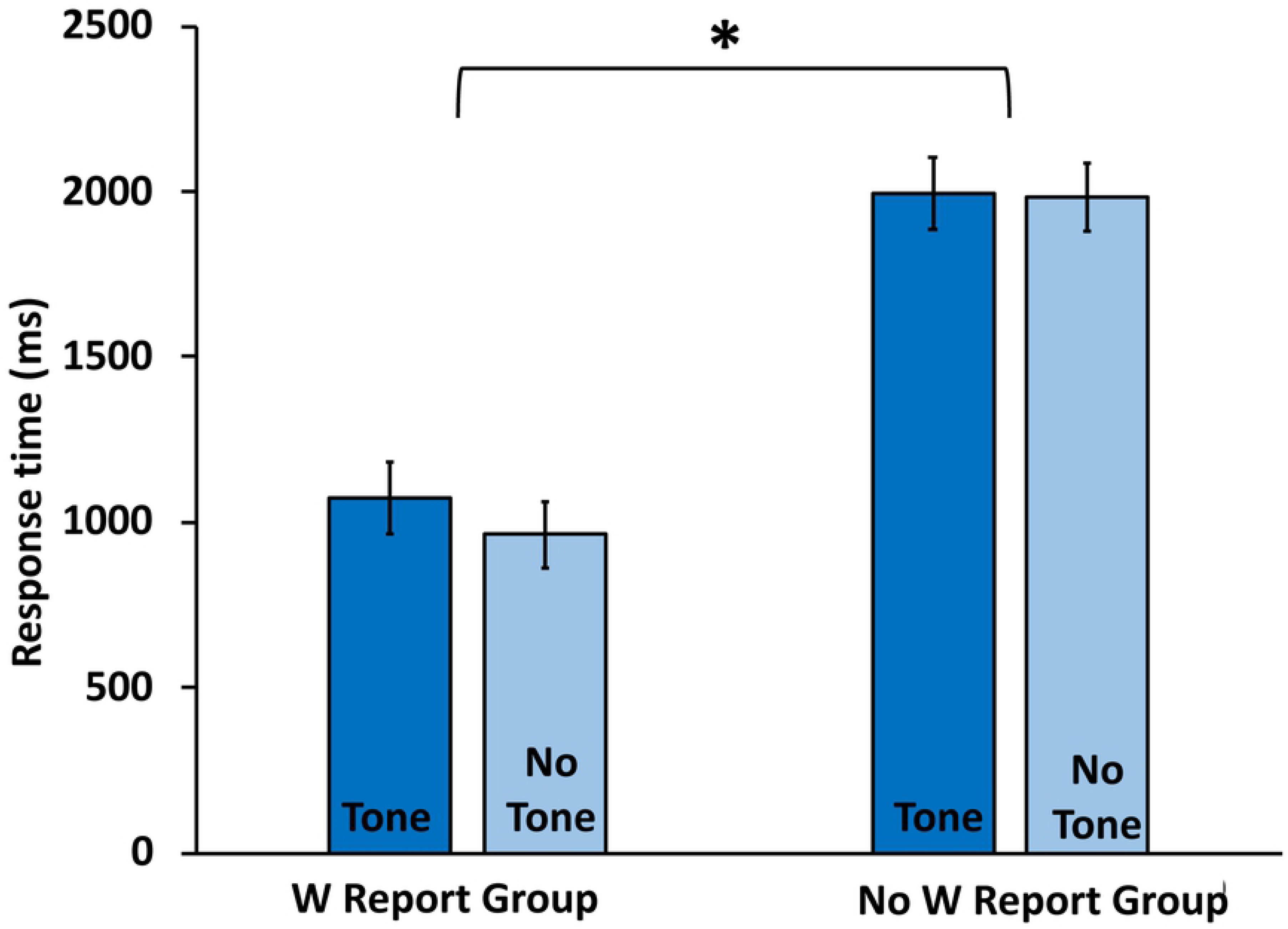
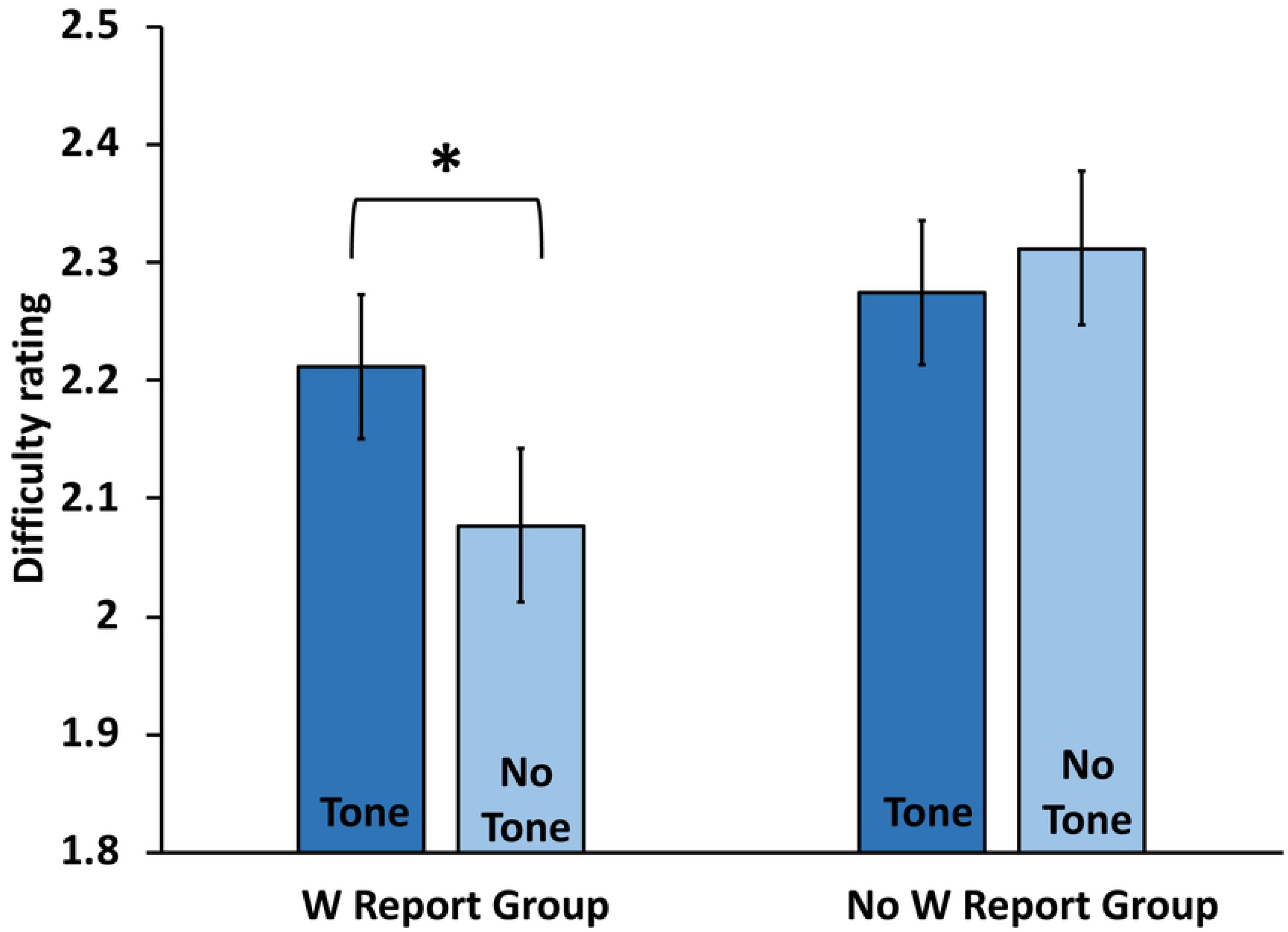
Reported W time, response time, and difficult rating as a function of Tone Presence and W Report Group. Analyses revealed W and response time dissociated such that W, but not response time, varied with Tone Presence. Difficulty varied with Tone Presence, but only when W was also reported, suggesting that the W experience had a role in difficulty assessment. Figure 2A: Reported decision time (W) as a function of Tone Presence. W was judged as later when the auditory feedback was delayed. Figure 2B: Response time as a function of Tone Presence and W Report Group. Response time did not vary with the tone manipulation. However, longer response time was observed in the No W Report group. Figure 2C: Difficulty rating as a function of Tone Presence and W Report Group. The tone manipulation affected the difficulty assessment such that the tone elicited a higher difficult rating. The effect of the tone manipulation was observed in the W Report Group but not in the No W Report Group.

##### Response time

Of the 24 participants recruited for the No W Report group, two participants’ response time values exceeded two standard deviations of the group mean. Therefore, only 22 participants in the No W Report group were included in response time analyses.

In contrast to W, we did not anticipate the tone manipulation to have an effect on response time. The response time data were subjected to a 2 Tone Presence x 2 W Report Group mixed ANOVA. Mean response time data are shown in Figure 2B. As predicted, the results revealed no main effect of Tone Presence, *F*(1,44) = 2.209, *p* = .144, η^2^ = .048, nor an interaction effect between the Tone Presence and W Report Group, *F*(1,44) = 1.822, *p* = .184, η^2^ = .040, suggesting that the tone manipulation did not have an impact on response time. These response time patterns differ from those for W and thus provide a condition in which response time and W are dissociated. In other words, while the presence of the tone had an effect on participant reports of W, it did not impact response time. Given this dissociation, we next examined how difficulty rating data were distributed in this condition.

There was a main effect of W Report Group on response time. The analysis revealed that response time was shorter in the W Report group (*M* = 1059.45, *SE* = 80.04) compared to the No W Report group (*M* = 2031.67, *SE* = 113.72), *F*(1,44) = 50.021, *p* < .001, η^2^ = .532, corrected for multiple comparisons. While not the primary focus of the current experiment, this finding is explored in the Discussion below.

##### Difficulty rating

The two previous analyses showed that W varied with the presence of the tone while response time did not. We next examined whether difficulty rating behaved in the same manner as W with respect to the presence of the tone. The difficulty rating was subjected to a 2 Tone Presence x 2 W Report Group mixed ANOVA. The analysis revealed an interaction effect between Tone Presence and W Report Group, *F*(1,46) = 5.105, *p* = .029, η^2^ = .100. Pairwise comparisons further showed that the interaction effect was attributed to the influence of Tone Presence in the W Report Group such that the difficulty rating was higher when the delayed tone was present (*M* = 2.21, *SE* = .06) than when the tone was absent (*M* = 2.08, *SE* = .07), *t*(23) = 2.45, *p* = .023, corrected for multiple comparisons, BF_10_ = 2.5. The difficulty rating scores in the delayed tone and no tone conditions, however, were not statistically different in the No W Report Group (*M* = 2.28, *SE* = .06 in the Delayed Tone condition; *M* = 2.31, *SE* = .07 in the No Tone Condition), *t*(23) = 0.71, *p* = .483, BF_01_ = 2.8. The difficulty rating data are represented in Figure 2C.

The interaction effect observed in this analysis suggests that when time W is explicitly accessed (i.e., W Report Group), the difficulty ratings are affected by the tone manipulation. However, the difficulty ratings are unaffected in the absence of W report. This key finding suggests that the temporal experience of W plays an important role in difficulty judgments.

In addition to the interaction effect, there was a main effect of the W Report. The ANOVA revealed that the W Report group’s mean difficulty rating (*M* = 2.15, *SE* = .06) was lower than the mean rating given by the No W Report group (*M* = 2.29, *SE* = .06), *F*(1,43) = 4.641, *p* = .037, η^2^ = .097. However, this did not pass the Benjamini-Hochberg test (BH *p*-value = .09) . Since the W Report variable was a between-subjects variable, it might be the case that there was greater variability in the sampled population, making the results inconclusive.

### Discussion

In this experiment, we have established a stance in which W and response time are dissociated: W, but not response time, varies with a delayed auditory feedback. The observed main effect of the tone manipulation on W was expected and served as a manipulation check. The absence of a main effect of the tone manipulation on response time was also expected since the tone came after the motor response and therefore should not have an influence on the timing of the motor execution.

Our key finding is the effect of the tone manipulation on difficulty assessment. We have demonstrated that when the tone was presented, difficulty ratings were higher than when the tone was absent. Importantly, this effect was observed only in the W Report group, but not in the No W Report group. An interpretation of this key interaction effect is that the tone manipulation influenced W in the W Report group, and in turn, W influenced difficulty ratings. In the absence of the W report, the tone manipulation was ineffective. This interpretation therefore implies that the temporal experience of W plays a role in subsequent metacognitive process of difficulty assessment.

The post-interview results observed in the No W Report group further emphasize the influence of W as a conscious subjective experience. The interview provided additional qualitative data that participants in the No W Report group were not monitoring their temporal experience and did not explicitly incorporate timing information. As a result of inattentiveness to timing of decision, difficulty ratings did not vary with the tone in the No W Report group. Aligned with the AIR theory (Prinz, 2005), this observation is consistent with the perspective that timing of decision may possibly reside at a lower level of consciousness (Mylopoulous, 2015). When instructed to explicitly attend to it, the timing of decision then emerges to a more aware, intermediate level of consciousness. Upon entering this stage, the timing information consequentially affects the difficulty assessment.

A concern with the results is that the change in difficulty rating might not have been related to the timing of decision, but instead is an artifact related to an interruption in the performance due to the presence of the tone. This concern can be addressed by the interaction effect observed in the difficulty rating analysis. Recall that the tone manipulation was administered to both the W Report and the No W Report groups. The effect of the tone manipulation on ratings, however, was observed only when the W report was solicited. Therefore, this suggests that interference of the tone, if any, was not the primary contribution of the tone. Rather, it was the W experience that played a significant role in difficulty assessment.

It is also important to note the main effect of the W Report on response time; shorter response time was observed in the W Report group compared to the No W Report group. One speculation is that the participants in the W Report group might have been more attentionally engaged in the overall task given they were explicitly asked to identify and remember W. Given this focus, it could have led to a faster response time. If this speculation were empirically supported, it would add to the claim that awareness of the reported time of decision (has consequences in related thoughts and action.

The results thus far provide evidence for a close relationship between subjective timing of decision (W) and difficulty ratings. However, it is unclear if the relationship is unidirectional such that W affects difficulty rating, or whether anticipated difficulty level has an impact on W, as it could be speculated that W is inferred from the ratings. We examined these concerns in Experiment 2.

## EXPERIMENT 2

In Experiment 2, we investigated whether prospective knowledge or perception of difficulty influences the perceived time W. To manipulate the perception of difficulty, two sets of stimuli of medium-rated difficulty were assigned to either an EASY or DIFFICULT set as detailed below. W reports for these “manipulated easy” and “manipulated difficult” stimuli were compared. If the manipulation influenced the W report when evaluating these stimuli, it would imply that W and difficulty judgment covary. On the other hand, if W did not vary with the difficulty manipulation, it would suggest that the relationship observed in Experiment 1 is unidirectional.

### Methods

#### Participants

Twenty-four participants (17 females, 18-50 years old) were recruited from a pool of undergraduate student volunteers at the University of California, Davis. The stimuli were drawn based on survey results given by separate, independent participants (67 participants; 44 females, 18-32 years old) also recruited from a pool of undergraduate volunteers at the University of California, Davis. All participants were fluent in English. All participants consented in writing to the study protocol which followed the guidelines approved by the Institutional Review Board of the University of California, Davis.

#### Rationale and estimation of sample size

Consistent with Experiment 1, we recruited 24 participants.

#### Survey and Stimuli

A separate group of participants (N = 67) rated the difficulty of making an Agree/Disagree judgment for each of 111 statements using a Likert scale of 1 to 5 (5 being most difficult). The 15 easiest and the 15 most difficult statements were assigned to the “True Easy” and “True Difficult” conditions, respectively. We next identified 30 statements that were rated of medium difficulty (ranked between 41 to 70 out of the 111 statements). These 30 statements were further divided into two sets of stimuli. To ensure the overall equality of the two sets, statements were assigned to each set according to their ranking order being odd or even; odd ranking numbers were assigned to Set A, and even ranking numbers were assigned to Set B. The 15 True Easy statements and the 15 statements from Set A (Manipulated Easy) made up the EASY condition. The set of 15 True Difficult statements and the 15 statements from Set B (Manipulated Difficult) made up the DIFFICULT condition. Assignment of the two medium sets to the EASY and DIFFICULT conditions was counterbalanced, and the 30 stimuli within each condition were randomized across participants. All stimuli for Experiment 2 can be found in Appendix 1.

#### Procedure

Participants were presented with a practice session (10 trials) followed by blocks of either 30 EASY or DIFFICULT statements, and then the remaining block of 30 DIFFICULT or EASY statements. The presentation order of EASY and DIFFICULT blocks was counterbalanced. As in Experiment 1, participants were asked to make a decision about each statement and to report W and the difficulty rating. To manipulate the perception of difficulty, we implemented the following scripts at the beginning of the EASY and DIFFICULT experimental blocks.

<u>EASY:</u> *From a survey of 100 people, these statements were considered to be easier to evaluate. On a scale of 1 to 5, 1 being very easy and 5 being difficult, the average rating was 1.72. We are curious how you would rate them.*
<u>DIFFICULT:</u> *From a survey of 100* people, *these statements were considered to be more difficult to evaluate. On a scale of 1 to 5, 1 being very easy and 5 being difficult, the average rating was 4.36. We are curious how you would rate them.*

Aside from these additional scripts, the remaining procedure and instructions were similar to the W Report (tone absent) condition in Experiment 1. Participants verbally reported W and provided a difficulty rating on a scale of 1 to 5 (5 being most difficult).

### Results

Three dependent variables, response time, difficulty rating, and W were subjected to separate within subject one-way ANOVAs and post-hoc pairwise comparisons. Two of the 24 recruited participants were not able to complete the task; therefore, their data were excluded from the analyses.

#### Difficulty rating

The primary purpose of this analysis was to demonstrate the effectiveness of the difficulty manipulation. We observed a significant difference across the four categories of stimuli, *F*(3, 63) = 68.495, *p* < .001, η^2^ = .765. Post-hoc pairwise comparisons showed that the True Easy condition was judged to be the easiest (*M* = 1.45, *SE* = .08), and the perceived difficulty significantly increased with Manipulated Easy (*M* = 1.90, *SE* = .12), Manipulated Difficult (*M* = 2.46, *SE* = .10), and True Difficult (*M* = 3.00, *SE* = .13), *p* < .001. Importantly, the difference in the ratings between the Manipulated Easy trials and the Manipulated Difficult trials was .56, and this was statistically significant, *t*(21) = 5.266, *p* < .001, corrected for multiple comparisons, BF_10_ = 762.2. The results thus served as a check that the manipulation of difficulty was effective. The difficulty rating data are depicted in Figure 3A. We next examined whether W varied according to the <u>manipulated</u> difficulty.

**Figure 3.**
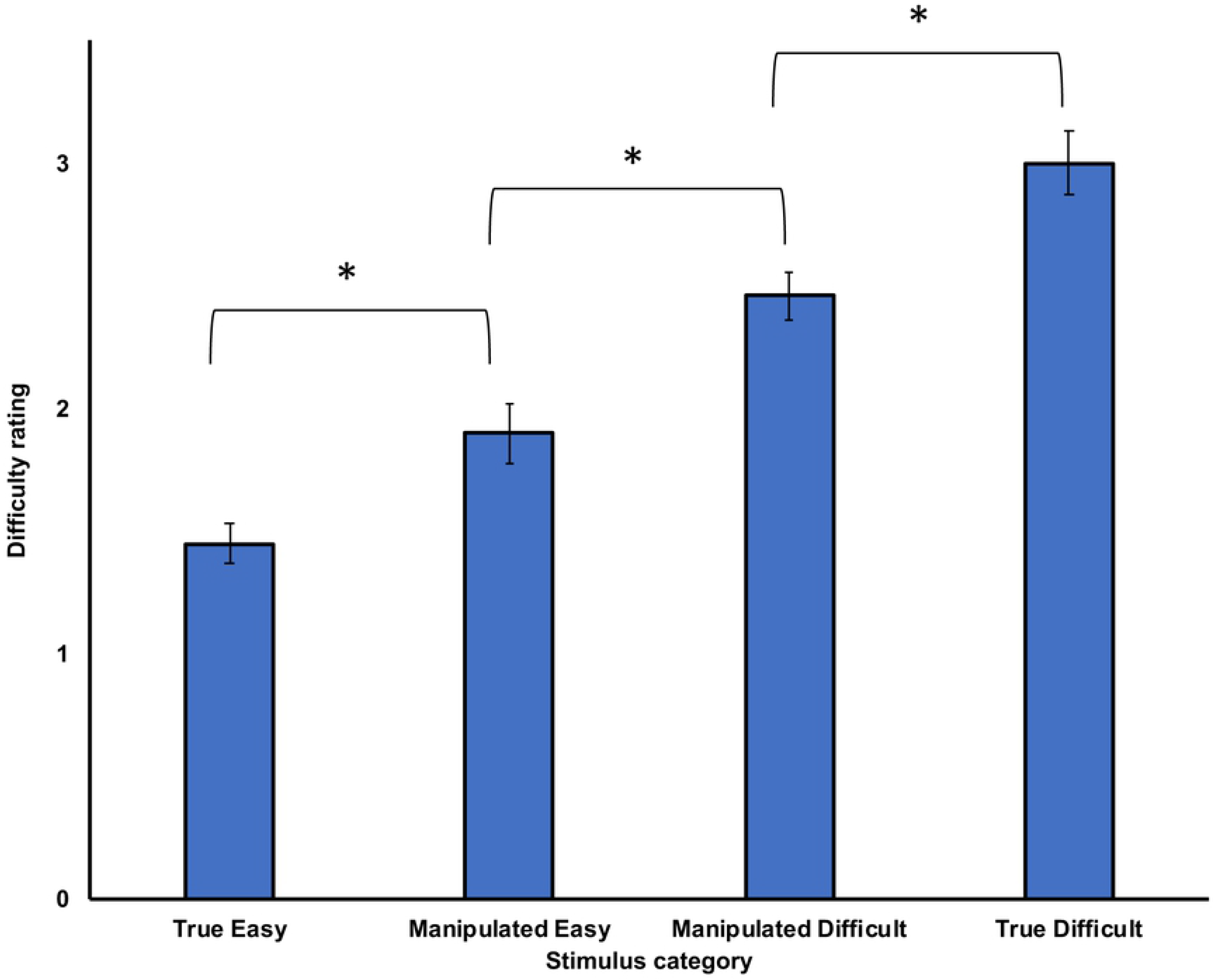
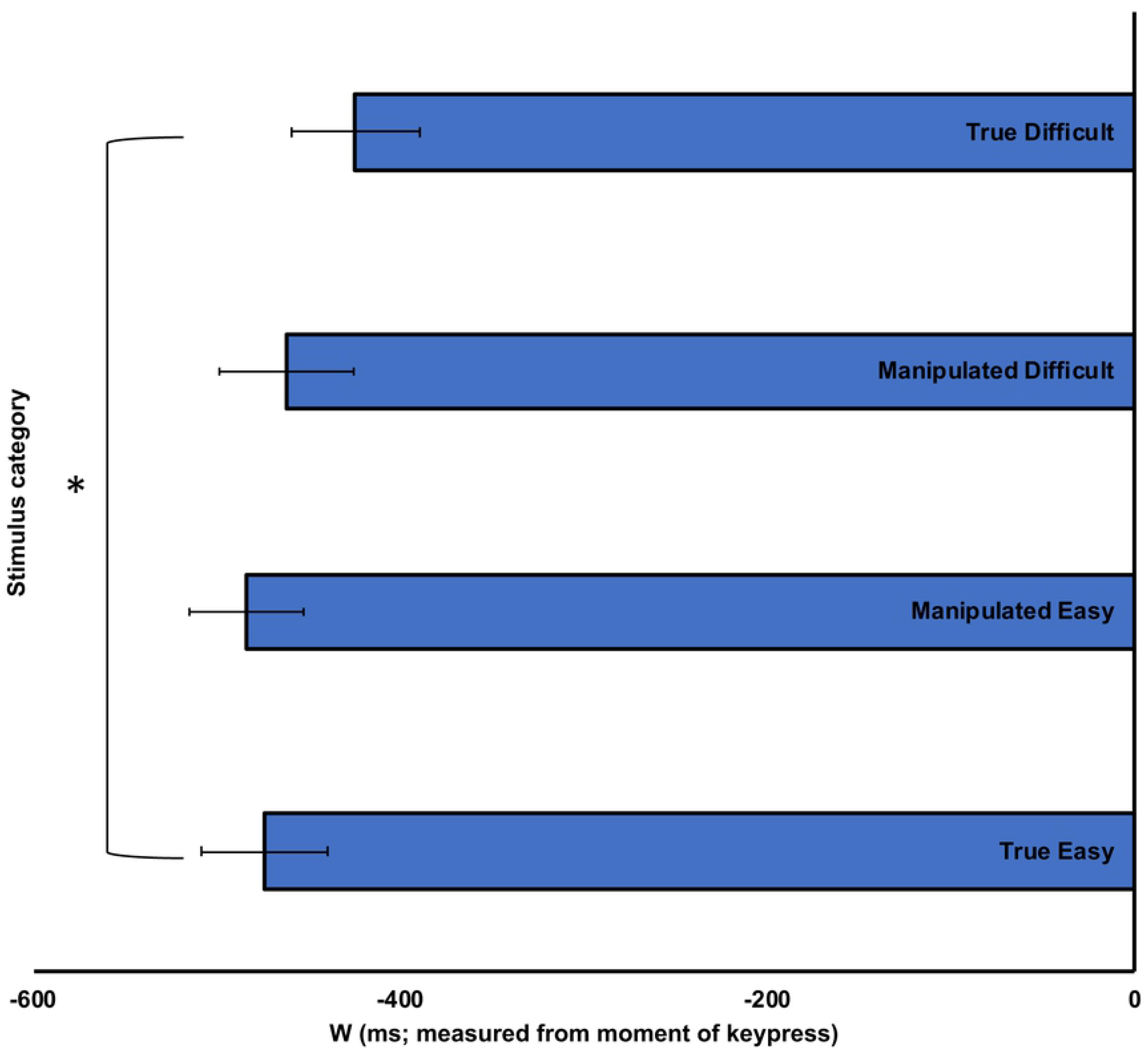
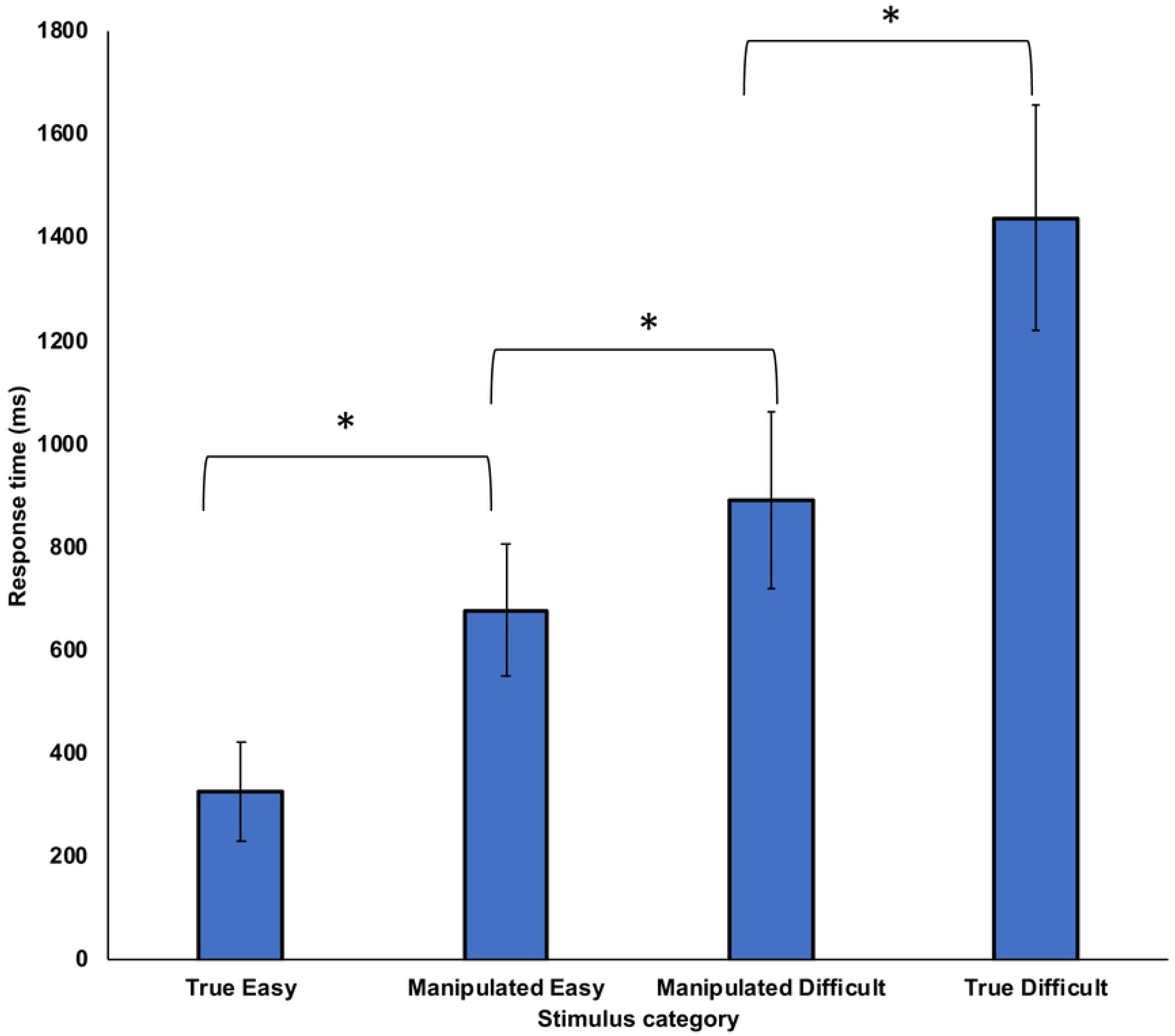
Difficulty rating, reported W and response time as a function of tone manipulation. Difficulty rating and response time varied with the difficulty manipulation (A and C), however, W did not (B). Figure 3A. Difficulty rating as a function of the difficulty manipulation. Medium-rated statement stimuli were placed in either the Manipulated Easy and Manipulated Difficult conditions. Lower difficulty ratings were given to these stimuli when placed in Manipulated Easy condition compared to the Manipulated Difficult condition. Figure 3B. Reported decision time (W) as a function of difficulty manipulation. Medium-rated stimuli were placed in the Manipulated Easy and Manipulated Difficult conditions. W reports were not statistically different between these conditions. Figure 3C. Response time as a function of difficulty manipulation. Medium-rated stimuli were placed in the Manipulated Easy and Manipulated Difficult conditions. Response time in the Manipulated Easy condition was shorter than in the Manipulated Difficult.

#### W

A one-way ANOVA showed a significant difference across the four categories of stimuli, *F*(3,63) = 3.699, *p* = .016, η^2^ = .150. Post-hoc pairwise comparisons further showed a significant difference in the W reports between the True Easy (*M* = 473.93 before keypress, *SE* = 34.51) and the True Difficult (*M* = 424.44 before keypress, *SE* = 35.01), *t*(21) = 2.150, *p* = .043, BF_10_ = 1.5, BH *p-*value = .052. Consistent with our *a priori* hypothesis, this finding replicated our prior report that an easy decision is associated with an earlier W and a difficult decision is associated with a later W (Isham et al., 2017). The critical observation though is the lack of statistical difference in the W reports between the Manipulated Easy (*M* = 483.54 before keypress, *SE* = 31.01) and Manipulated Difficult conditions (*M* = 461.95 before keypress, *SE* = 36.16), *t*(21) = 1.088, *p* = .289, BF_01_ = 2.7. This result suggests that difficulty judgments do not influence W. The W data are depicted in Figure 3B.

#### Response time

The response time was subjected to a one-way ANOVA. There was a main effect of the different types of trials, *F*(3,63) = 34.476, *p* = .000, η^2^ = .621. Post-hoc pairwise comparisons showed that the mean response time varied across the four conditions tested: True Easy condition (*M* = 326.47, SE = 94.85), Manipulated Easy (*M* = 677.61, *SE* = 127.32), the Manipulated Difficult (*M* = 890.26, *SE* = 169.89), and the True Difficult (*M* = 1436.44, *SE* = 218.31), *p* < .023. Although not the primary focus of the study, the results showed a significant difference of 212.64 ms between the mean response time of the Manipulated Easy and Manipulated Difficult, *t*(21) = 2.447, *p* = .023, corrected for multiple comparisons, BF_10_ = 2.5. This suggests the possibility that prospective knowledge about difficulty may influence planning on how much time to spend on a decision. The response time data are depicted in Figure 3C.

### Discussion

The purpose of Experiment 2 was to examine whether biases in the perception of difficulty influence the estimates of W or response time. We successfully biased the belief that the target stimuli, which otherwise were equated, were either easy or difficult to evaluate. Despite the effectiveness of the manipulation, the manipulated stimuli did not influence W. In conjunction with the results from Experiment 1, this lack of effect on W supports the perspective that the relationship between W and difficulty is unidirectional such that W influences the perceived degree of difficulty, but not the reverse.

With respect to response time, the results suggest the possibility that response time covaries with perceived difficulty. It might be the case that a trial, when perceived as easy, encourages participants to spend less time making the decision; and a trial, when perceived to be difficult, encourages participants to spend more time deliberating before reaching a decision. This corroborates a previous observation that perceived difficulty influences response time (e.g., Kukla, 1974); however, the result is inconsistent with the view that response time contributes to the experience of task difficulty (e.g., Descender, Opstal, & Van den Bussche, 2017).

## General Discussion

Brass and Haggard (2008) propose the temporal concept of “When” as one of the three key components underlying decision making. The current study investigates whether the subjective temporal experience of decision (W) modifies the metacognition of difficulty judgments. In two experiments, we observed that the temporal experience of decision had the priority over actual response time in modifying judgments regarding decision difficulty. That is, in Experiment 1, we observed that the perception of difficulty varied with the perception of when a decision was made (time W). When participants experienced delayed auditory feedback, they judged W to be later and reported their decisions to be more difficult compared to when there was no tone and they judged W to be earlier. On the other hand, when we manipulated perceived difficulty, participants did not change their reported time of W (Experiment 2).

A possible explanation to this observation is built on the notion that decision making operates within a hierarchical framework. This framework begins with the thought of the decision, whose timing is indexed by W, that precedes the evaluation of decisional difficulty (difficulty rating). This, in turn, precedes the total time (response time) spent on the decision-making task. It would be quite natural to form a perception of decisional difficulty in terms of when we think the decision was made. In contrast, it is less natural, although not implausible, to first think that the decision is difficult, and to then spend a longer time deliberating on it.

The idea of hierarchical order of events is further supported by the findings from Experiment 2, which showed that the manipulated difficulty impacted response time. This effect may be related to planning; when we expect a decision to be difficult, we may plan to spend longer time on the problem. Further investigation is necessary to verify the possibility of this causal relationship. If supported, this would challenge the perspective that response time influences difficulty ratings (e.g., Desender et al., 2017).

### Mechanisms related to the function of consciousness

We propose that the observed effect of W on the difficulty ratings leads to the interpretation that consciousness serves to modify future thoughts and behaviors. While we believe the current findings support this perspective, the mechanistic component related to our findings remains unresolved. As discussed earlier, our results may map on to the framework of the AIR theory offered by Prinz (2000). According to this theory, consciousness serves the function of sending viewpoint specific information from a lower sensory level to an intermediate level of processing. Even so, it is unclear what facilitates the migration from one level to the next.

A memory-based theory of consciousness may help explain how this is possible. Evidence summarized by Kanai et al. (2019) suggests the possibility that consciousness is involved in maintaining sensory information in short term memory. Such an idea connects with the findings from our current study. In our paradigm, participants maintained and integrated sensory information from visual (clock reading), tactile (button press) and auditory (feedback tone) modalities to produce a W report. Linking this to the AIR theory, we speculate that consciousness’ role in short term memory maintenance helps keep the integrated sensory information at the intermediate level or higher. In so doing, it is possible that the intermediate level is also where meta-cognition about the timing of a decision is formed or updated. Generally speaking, memory is flexible, fluid, and is likely updated when new information is available (e.g., Loftus & Palmer (1974)’s landmark study showed how additional information can alter one’s memory of a witnessed car accident) . We believe this may generalize to memory for the timing of decision; when presented with a tone, participant’s perception of W is updated. Subsequently, when prompted to evaluate difficulty, we infer from this most current version of W to produce the difficulty report.

Another key finding from the current study is that the effect of W on difficulty judgment was only observed when W was explicitly attended. This observation, and in conjunction with the fact that the experimental task involved non-perceptual, subjective statement stimuli, implies that the reported W was a more deliberate and of a higher-level conscious thought process. This conclusion concurs with a previous study whose drift diffusion approach illustrated that decision time was a deliberative process, rather than perceptual (Kang, Petzschner, Wolport & Shadlen (2017). Upon this assumption, the current findings have the potential of addressing the extent in which conscious and non-conscious decision time play a role in the metacognition of difficulty judgment.

Related, while it is beyond the scope of the current study to specify the mechanisms in which W affects difficulty ratings, one speculation resonates with the idea that the effect of W on difficulty ratings operates as a low-level priming effect. That is, an early W could prime an easy rating and a late W could prime a higher difficulty rating. If additional investigation demonstrated support for such low-level priming effect, it would invite additional questions surrounding W and difficulty ratings. Particularly, why would low-level priming effect apply only in the context in which the W unidirectionally affected ratings (i.e., Experiment 1), but presumably would not apply in the reverse scenario (i.e., difficulty judgment did not affect W judgment, Experiment 2). This context-dependent intersection between metacognition and low-level cognition thus could expand further the relationship between consciousness and decision making.

### Limitations

The experimental manipulation in Experiment 2 was used to bias the perception of difficulty of previously rated medium-difficulty stimuli. To best maintain the integrity of the manipulation, we could not assess for baseline ratings from the target participants; we could only assume that the focus participants had a similar perception of difficulty as the independent raters.

### Future directions

In addition to further investigation to address some of the speculations described above, our results may also inspire other research related to temporal consciousness.

#### Intentional Binding

The evidence from Experiment 2 implying that difficulty rating does not influence W questions the robustness of Intentional Binding theory. According to the theory, an effortful action leads to a temporal compression between the timing of action and the ensuing effect (Haggard, Clark & Kalogeras, 2002). Results from Experiment 2, however, showed that perceiving a decision to be effortful and difficult led to a time dilation rather than a time compression between W and keypress. Our results provide an instance where Intentional Binding may not apply. Future studies may wish to explore this in depth by acquiring the traditional measures used in the Intentional Binding paradigm.

#### Applications

A recent study showed that altering perceived time also influenced physiological responses (e.g., perceived time alters the blood sugar level in people with type 2 diabetes; Park, Pagnini, Reece, Phillips & Langer, 2016). Their results provide additional support to the perspective that temporal experience plays an influential role beyond behavioral modifications. Collectively, our results and Park et al.’s observations may provide a basis for future applications that utilize time perception to better our physical, mental, and cognitive functions.

## Conclusions

In summary, the current study has provided additional insights into Libet’s W. As an index of temporal consciousness, W has been controversial. However, despite the mechanistic questions surrounding W, here we have shown that this subjective temporal experience plays a significant role in modifying subsequent thoughts and behaviors related to the assessment of task difficulty.

## Acknowledgement

The author thanks Mary Peterson, Melinda Davis, Arne Ekstrom, Dale Berger, and Sara Lomayesva, along with the previous anonymous reviewers, for valuable comments on an earlier version of the manuscript. The author also thanks The Evaluation Group for Analysis of Data (EGAD) for assisting in the analyses. The author also thanks Bevy John, Jon Hui, and Daryan Singer for their assistance with data collection. The study was partially supported by research award from UC Davis Center for Effective Teaching and Learning.

## Appendix 1

<u>True Easy</u> –15 stimuli with lowest mean difficulty rating

I would give up a friend for money.

I would give up a friend for popularity.

I am happy for the success of my friends.

I support women’s rights.

I support children’s rights.

I would go to jail for a night of fun.

I would go to jail for money.

It’s okay to drive under the influence if you feel fine.

It’s okay to drive under the influence if you are hungry.

It is fine to steal from your best friend’s cousin.

It is fine to steal from your best friend’s sibling.

I want others to be respectful to me.

I want others to worship me.

I would give up my savings to pay for my mother’s hospital bills.

I believe in fate.

<u>Set A</u> – 15 stimuli with moderate mean difficulty rating

Money can’t buy happiness.

To save the village it is okay to sacrifice a child.

I would give up my savings to pay for my mother’s new car.

Children should not be allowed access on Facebook.

If I found money, I would keep the cash.

You should not curse in front of women.

It is okay to speed when I am running late to class.

It is okay to speed when there is a family emergency.

It is okay to have sex before marriage.

The strength of a nation is judged by the way its people are treated.

In a desperate situation, it is okay to consume a sick child for food.

In a desperate situation, it is okay to consume a dog for food.

I would give up friends for more friends.

I am happy for the success of strangers.

Bad people deserve to be punished.

<u>Set B</u> – 15 stimuli with moderate mean difficulty rating

I would do anything for a million dollars.

Freedom is more important than money.

It’s okay to cheat in a game.

It’s okay to publically criticize celebrities.

Children should not have cell phones.

It is my responsibility to help conserve energy.

It is my responsibility to help conserve wildlife.

If I found money, I would return it.

A cold-blooded killer should be tortured.

It is better to look forward than to look back.

It is better to build a career before marriage.

If I saw someone being attacked, I would help.

I sympathize with wealthy people who have just lost their jobs.

okay to bend the rules to help someone out.

It is fine to steal from your best friend’s enemy.

<u>True Difficult</u> – 15 stimuli with highest mean difficulty rating

Lying is okay if it keeps me safe.

Lying is okay if it keeps my country safe.

I am happy for the success of my enemies.

I sympathize with wealthy people who have no real friends.

If I won the lottery, I would give it all to my family.

A person’s true character is revealed through power.

A person’s true character is revealed through love.

A person’s true character is revealed through leadership.

Freedom is more important than death.

Freedom is more important than love.

I would steal to give to my children.

It’s better to forget than to forgive.

It’s better to remember pain than to forget pleasure.

It’s better to co-habitat before marriage.

I’d rather receive an A− than B+.

## References

Banks W, Isham EA. We infer rather than perceive the moment we decided to act. Psychological Science. 2009; 20: 17–21.

Baumeister RF, Masicampo EJ, Vohs KD. Do conscious thoughts cause behavior?. Annual review of psychology. 2011; 62: 331–361.

Benjamini Y, Drai, D, Elmer G, Kafkafi N, Golani I. Controlling the false discovery rate in behavior genetics research. Behavioral and Brain Research, 2001; 125: 279–284.

Brass M, Haggard P. The what, when, whether of intentional action. Neuroscientist. 2008; 14: 319–325.

Desender K, Van Opstal F, Van den Bussche E. Subjective experience of difficulty depends on multiple cues. Scientific Reports, 2007; 7, 44222, doi: 10.1038/srep44222

Eagleman DM, Sejnowski TJ. Motion signals bias localization judgments: A unified explanation for the flash-lab, flash-drag, flash-jump, and Frohlich illusions. Journal of Vision. 2007; 7, doi: https://doi.org/10.1167/7.4.3

Faul F, Erdfelder E, Lang A, Buchner A. G*Power 3: A flexible statistical power analysis program for the social, behavioral, and biomedical sciences. Behavior Research Methods. 2007: 39; 175–191.

Fleming SM, Massoni S, Gajdos T, Vergnaud, JC. Metacognition about the past and future: Quantifying common and distinct influences on prospective and retrospective judgments of self-performance. Neuroscience of Consciousness. 2016; 1. DOI: https://doi.org/10.1093/nc/niw018

Haggard P, Clark S, Kalogeras J. Voluntary action and conscious awareness. Nature Neuroscience. 2002; 382–385.

Isham EA, Wulf KA, Mejia C & Krisst LC. Deliberation period during easy and difficult decisions: re-examining Libet’s “veto” window in a more ecologically valid framework. Neuroscience of Consciousness. 2017; 3(1), nix002.

Loftus EF, Palmer JC. Reconstruction of automobile destruction: An example of the interaction between language and memory. Journal of Verbal Learning & Verbal Behavior. 1974; 13(5): 585–589.

Kang YHR, Petzschner FH, Wolpert, DM, Shadlen, MN. Piercing of consciousness as a threshold-crossing operation. Current Biology. 2017; 27: 2285–2295.

Kanai R, Chang A, Yu Y, de Abril IM, Biehl M, Guttenberg N. Information generation as a functional basis of consciousness. Neuroscience of Consciousness. 2019; doi: 10.1093/nc/niz016.

Kornhuber HH, Deecke, L. Changes in the brain potential in voluntary movements and passive movements in man: readiness potential and reafferent potentials. Pflugers Archiv fur die gesamte Physiologie des Menschen und der Tiere. 1965; 284: 1–17.

Kukla A. Performance as a function of resultant achievement motivation (perceived ability) and perceived difficulty. Journal of Research in Personality, 1974; 4: 374–383.

Libet B. Unconscious cerebral initiative and the role of conscious will in voluntary action. Behavioral and brain sciences. 1985; 8(4): 529–539.

Libet B. Do we have free will? In: Sinnot-Armstrong W, Nadel L, editors. Conscious will and responsibility. New York: Oxford University Press; 2011. p. 1–10.

Libet B, Gleason CA, Wright EW, Peal DK. Time of conscious intention to act in relation to onset of cerebral activity (readiness-potential); the unconscious initiation of a freely voluntary act. Brain. 1983; 106(3): 623–642.

Masicampo EJ, Baumeister RF. Conscious thought does not guide moment-to-moment actions—it serves social and cultural functions. Frontiers in Psychology. 2013; 4: 478. doi: 10.3389/fpsyg.2013.00478

Mylopoulous M. Conscious intention: A challenge for AIR theory. Frontiers in Psychology. 2015; 6: 675. doi:10.3389/fpsyg.2015.00675

Park C, Pagnini F, Reece A, Phillips D, Langer E. Blood sugar level follows perceived time rather than actual time in people with type 2 diabetes. Proceedings of the National Academy of Sciences. 2016; 19: 8168–70.

Pham LB, Taylor SE (1999). From thought to action: Effects of process-versus outcome-based mental simulations on performance. Personality and Social Psychology Bulletin. 1999; 25(2): 250–260.

Pockett, S. (2004). Does consciousness cause behavior? Journal of Consciousness Studies. 2004; 11(2): 23–40.

Prinz JJ. A neurofunctioal theory of visual consciousness. Consciousness & Cognition. 2000; 9: 243–259. 10.1006/ccog.2000.0442

Prinz JJ. A neural functional theory of consciousness. In Brooks A., Akins K, editors. Cognition and the Brain: The Philosophy and Neuroscience Movement. New York: Cambridge University Press; 2005. p. 381–396.

Rouder JN, Speckman PL, Sun D, Morey RD, Iverson G. Bayesian t-tests for accepting and rejecting the null hypothesis. Psychonomic Bulletin & Review. 2009; 16: 225–237.

Wegner D. The Illusion of Conscious Will. Cambridge MA: The MIT Press; 2002.

